# Improving N-Glycosylation and Biopharmaceutical Production Predictions Using AutoML-Built Residual Hybrid Models

**DOI:** 10.1101/2024.08.27.609988

**Authors:** Pedro Seber, Richard D. Braatz

## Abstract

N-glycosylation has many essential biological roles, and is important for biotherapeutics as it can affect drug efficacy, duration of effect, and toxicity. Its importance has motivated the development of mechanistic models for quantitatively predicting the distribution of N-glycans during therapeutic protein production. Here we present a residual hybrid modeling approach that integrates mechanistic modeling with machine learning to produce significantly more accurate predictions for production of monoclonal antibodies in batch, fed-batch, and perfusion cell culture. For the largest dataset, the residual hybrid models have an average 736-fold reduction in testing prediction error. Furthermore, the residual hybrid models have lower prediction errors than the mechanistic models for all of the predicted variables in the datasets. We provide the automatic machine learning software used in this work, allowing other researchers to reproduce this work and use our software for other tasks and datasets.

## 1 Introduction

Glycosylation is a protein co-translational and post-translational modification that involves adding a glycan or glycans to proteins. N-linked glycosylation, a subtype of glycosylation, occurs when a glycan is added to the nitrogen of an asparagine or arginine. N-glycosylation contributes to many essential functional and structural roles [1, 2, 3]. Highlighting the importance of N-glycosylation, improper glycosylation or deglycosylation is associated with multiple diseases, including cancers [4], infections [5], and congenital disorders [6].

There is great interest in N-glycosylation from the biomedical and pharmaceutical industry, physicians, and patients because of its high therapeutical and diagnostic relevance. In many types of carcinoma, increases in fucosylation, branching, and sialylation occur [7]. Disialoganglioside is expressed by almost all neuroblastomas, and Phase I–III studies have shown that anti-disialoganglioside monoclonal antibodies can be successful against those [8, 9]. Poly-*α*2,8-sialylation, for example, increases the half-lives of antibodies but does not lead to tolerance problems [10]. Conversely, the presence of glycans foreign to humans can be detrimental to a therapeutic. N-glycolylneuraminic acid is immunogenic to humans [11] but is present in some CHO-cell-derived glycoproteins [12].

Despite these critical functions of N-glycosylation in biotherapeutical contexts and multiple developments in this field, such as the genetic engineering of CHO cells to increase glycoprotein sialylation [13], some challenges persist. Proteins can be glycosylated in multiple locations, so any investigations need to elucidate not only the glycan compositions but also where each glycan is located [7]. This structural and regional diversity makes it challenging to determine specific functions of N-glycans [3]. In a diagnostic context, analyzing patient glycosylation samples is also challenging due to the complex equipment needed, which has limited the progress of personalized medicine [7].

To assist in better understanding and predicting N-glycosylation, many computational models have been created, which may be subdivided into mechanistic and data-driven models. Mechanistic models use physical knowledge, typically in the form of differential equations, to make predictions. They require little-to-no process data and always output answers that are constrained by physics; however, they also demand significant understanding of the system and can be slow [14], a problem alleviated by model simplifications [15, 16]. Data-driven models directly leverage experimental data to make predictions. They require zero domain-based knowledge (although they can benefit from it), are typically fast once trained, and the more complex data-driven models are universal function approximators [17, 18, 14]; however, they require high amounts of high-quality data due to their high variance, can produce non-physical outputs, and are typically not interpretable [14, 19]. Most works focused on N-glycosylation, particularly those focused on predicting the distribution of N-glycans, use mechanistic models due to a lack of high-quality data and data-driven modeling knowledge. The literature on mechanistic models for the prediction of N-glycan distributions is extensive; some examples can be found in the reviews of Refs. [20] and [21]. Significant works using data-driven models for the same task include Refs. [22] and [23]. An alternative to mechanistic and data-driven models are hybrid models. Hybrid models seek to combine these two types of models to create something with the advantages of both and the disadvantages of neither [14]. Although of high interest, the construction of hybrid models is not straightforward. First, they require knowledge of both mechanistic and data-driven modeling to be successfully implemented. Second, hybrid learning encompasses many architectures and model integration methods, and there have not been systematic studies on how to determine the best hybrid learning method for a given problem or system. Multiple examples of these architectures can be found in Refs. [14, 24, 25]. One of those architectures is the Residual Hybrid Model,^1^ in which a data-driven model learns the residuals (prediction errors) of a mechanistic model [26]. Despite the simplicity of this hybrid architecture, it has found success in many scientific problems [26, 27, 24, 14]. An illustration of this architecture is available in Fig. 1; other illustrations are also available in Fig. 1-A1 of Ref. [24] and Fig. 3 of Ref. [14].

**Figure 1:**
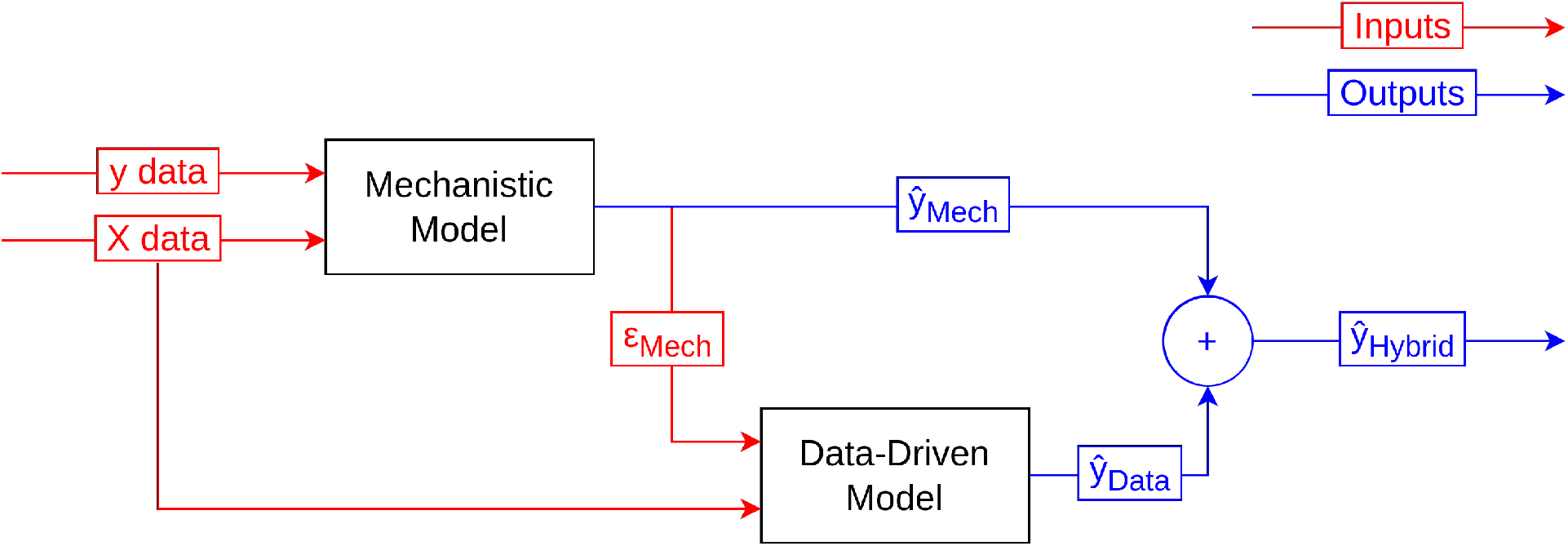
The residual hybrid model architecture. *ε*_Mech_ refers to the residuals (errors) of the predictions made by the mechanistic model. ŷ refers to the predictions made by a certain model architecture.

In this work, we construct residual hybrid models by combining mechanistic models from the literature with Lasso-Clip-EN (LCEN) [28] and artificial neural network (ANN) models trained by us. The data-driven models are trained using an efficient automatic machine learning (AutoML) software developed by us that completely eliminates the need for the end-user to have any knowledge in data-driven modeling. This software is free and open-source. These hybrid models are first used to predict the distribution of N-glycans attached to antibodies produced by Chinese hamster ovary (CHO) cells under different culture conditions. Then, the models are used to predict metrics that are relevant to biopharmaceutical production in CHO cells, including the titer, the galactosylation index of the products, and the levels of different chemicals in the culture medium over time. These CHO cultures were done in perfusion, fedbatch, and batch bioreactors depending on the dataset. Our hybrid models reduce the average prediction error by 149-fold on independent test sets when compared to the mechanistic models, and always lead to a reduction in the average test-set prediction error on the datasets investigated in this work. For the largest dataset, residual hybrid models have a test-set error that is 736-fold smaller than that of the mechanistic model.

## 2 Materials and Methods

This section describes the datasets and the residual hybrid models training methods. Instructions on running our AutoML software or recreating this article’s results are provided in Supplementary Information and in our GitHub repo, available at https://github.com/PedroSeber/SmartProcessAnalytics.

### 2.1 Datasets

Four previously published works provide the data used to train the hybrid models in this work. The first dataset, from Ref. [29], comprises the levels of N-glycans on monoclonal antibodies produced in a CHO perfusion culture as a function of viable cell density and the concentration of galactose and manganese ions. The second, from Ref. [30], comprises the levels of N-glycans on monoclonal antibodies produced in a CHO fed-batch culture, but as a function of pH, galactose concentration, and manganese ion concentration under different conditions and feeding strategies. This dataset also includes a time-based component, and galactose or manganese supplementation at specific time points was performed in some samples. The third, from Ref. [31], comprises the titer and galactosylation index of monoclonal antibodies produced in CHO fed-batch culture as a function of feed galactose and uridine. Finally, the fourth, from Ref. [32], comprises the levels of different chemicals (such as ammonium, lactate, or aspartic acid) in the culture media over time. The culture was done in a batch bioreactor as detailed in Refs. [32] and [33].

Data are split between cross-validation (CV) and test sets. For the Ref. [29] dataset, the same split used in that work is used here, and 3-fold CV is used. For the Ref. [30] dataset, the same split used in that work is used here, and 4-fold CV is used. For the Ref. [31] dataset, points FS2, FS3, and FS7 are used as the test set; the first two because they are outside of the design spec set by Ref. [31], and the last because the model of Ref. [31] had the highest prediction error on that sample. 4-fold CV is also used. For the Ref. [32] dataset, all points in the death phase (t > 135 hrs) are used as the test set, and 5-fold timeseries CV is used. These choices of test sets avoid test set leakage by ensuring the test set is sufficiently different from the training/cross-validation set. For robustness, the CV procedure is repeated 10 times for each combination of hyperparameters.^1^

### 2.2 Data-driven and residual hybrid model creation

Ordinary least-squares (OLS), elastic net (EN), LCEN, and ANN models are directly trained on the data using our AutoML software to serve as a baseline (in addition to the respective mechanistic models). OLS and EN models are constructed with scikit-learn [34], LCEN models are constructed as per Ref. [28], and ANN models (specifically, multilayer perceptrons [MLPs] and recurrent neural networks [RNNs]) are constructed with PyTorch [35] within our AutoML software. Furthermore, LCEN and ANN models are trained on the residuals of the mechanistic models to create residual hybrid models. A list of the hyperparameters used for each model architecture is available in Section S2. The best combination of hyperparameters for each model and task is determined by grid search, and the combination with the lowest cross-validation average loss (averaged over 10 repeats) is selected. Errors on an independent test dataset are then reported.

## 3 Results

### 3.1 Residual hybrid models improve N-glycan distribution predictions

This section includes the results of the models trained with the datasets of Refs. [29] and [30]. Ref. [29] featured two forms of models: a mechanistic model that used differential equations (named “Mechanistic” in this work) and a response surface methodology (RSM) model with intercept, linear, and 2nd-degree interaction terms. These models provide good fits even on an independent test set (Figs. 6 and 7 of Ref. [29]), with the mechanistic model providing a better fit. Nevertheless, these models still had high errors for some non-minor glycan forms; for example, the mechanistic model had a 47.9% and a 9.31% relative error when predicting the amount of antibodies with high-mannose (HM) and FA2G2 glycosylation respectively (Table 1). As per Section 2.2, we trained data-driven models to expand the models serving as a baseline and an MLP-based residual hybrid model. The OLS and EN models are very similar to the RSM model, but they lack the interactions present in the RSM model. Despite that, they provided similar (slightly worse) performance, indicating that the contribution of the interaction terms is limited yet not insignificant. LCEN had higher prediction accuracy than RSM for all four glycans, whereas the MLP model achieved the same on 3/4 glycans and had a 3.9% worse performance on the remaining glycan (Table 1). These results indicate that nonlinear terms can be important if they are not binary interaction terms, which is corroborated by how LCEN frequently selected features of the form 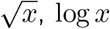, and 1*/*(*x*_*j*_*x*_*k*_). Nevertheless, with the exception of the MLP model to predict levels of HM, all of these data-driven models were inferior to the mechanistic model. It is likely that the chief reason for these higher percent relative errors (PREs) is the scarcity of data, as only 8 points are available for model training. Despite this shortage of data and the high accuracy of the mechanistic model for most glycans, a residual hybrid model composed of the mechanistic model followed by an MLP always achieved a lower PRE than the mechanistic model (Table 1). On average, the residual hybrid model led to 2.45-fold average reductions in the relative errors of the mechanistic model. These great results highlight how residual hybrid models are useful in predicting N-glycan distributions even when few data points have been collected and a strong mechanistic model is already in use. They also highlight the effectiveness of our AutoML method for training an MLP to succeed the mechanistic model in this hybrid architecture.

**Table 1:** Test-set percent relative errors (PREs) for different model architectures predicting the levels of each major N-glycan on the dataset of Ref. [29]. Models “Mechanistic” and “RSM” are from Ref. [29]; their PREs are obtained from the published data within. Models “OLS”, “EN”, “LCEN”, and “MLP” are data-driven models from this work used as baselines. Model “Mechanistic + MLP” is a residual hybrid model from this work. “Train Mean” is the mean of the training data. The lowest PREs are highlighted in bold.

| Architecture Name | High mannose | FA2G0 | FA2G1 | FA2G2 |
| --- | --- | --- | --- | --- |
| Mechanistic<br>(From [29]) | 47.9 | 2.18 | 2.42 | 9.31 |
| RSM<br>(From [29]) | 147 | 27.7 | 15.3 | 28.2 |
| OLS<br>(Baseline) | 67.5 | 38.9 | 23.3 | 29.3 |
| EN<br>(Baseline) | 71.2 | 35.9 | 17.2 | 28.8 |
| LCEN<br>(Baseline) | 53.4 | 25.5 | 10.9 | 23.8 |
| MLP<br>(Baseline) | 42.3 | 26.9 | 15.9 | 20.5 |
| Mechanistic + MLP<br>(This Work) | **32.8** | **1.15** | **0.76** | **2.86** |
| Train Mean<br>(From [29]) | 75.7 | 39.9 | 26.1 | 82.3 |

To further validate the ability of residual hybrid models to achieve higher accuracy than mechanistic models, a second work with this type of data was used. Ref. [30] included only a mechanistic model, again using differential equations and named “Mechanistic” in this work. Ref. [30]’s mechanistic model had worse fits than that of Ref. [29], as the former had low errors only for FA2G0, average errors for FA2G1, and high errors for the other two glycans (Table 2). Once again, we trained data-driven models to expand the models serving as a baseline and an MLP-based residual hybrid model. Despite the slightly increase in the amount of training data, the OLS and EN models performed poorly – their predictions were worse than predicting with the average of the training set. The only exception was the EN model trained to predict FA2G2 levels, which displayed a good performance, which even surpassed the mechanistic model. LCEN had mixed results: in 2/4 tasks, it surpassed the training mean, and in 2/4 tasks, it surpassed the mechanistic model. Once again, these results highlight the importance of nonlinearities to predict the distribution of N-glycans. The final data-driven model, an MLP model, had the best performance out of all models in this task. All of the MLP predictions surpassed the train mean and the mechanistic model, reaching test-set prediction errors 1.5-fold smaller on average than the mechanistic model. The

**Table 2:** Test-set percent relative errors (PRE) for different model architectures predicting the levels of each major N-glycan on the dataset of Ref. [30]. Model “Mechanistic” is from Ref. [30]; its PREs are obtained from the published data within. Models “OLS”, “EN”, “LCEN”, and “MLP” are data-driven models from this work used as baselines. Model “Mechanistic + MLP” is a residual hybrid model from this work. “Train Mean” is the mean of the training data. The lowest PREs are highlighted in bold.

| Architecture Name | High mannose | FA2G0 | FA2G1 | FA2G2 |
| --- | --- | --- | --- | --- |
| Mechanistic<br>(From [30]) | 41.4 | 9.47 | 17.0 | 39.1 |
| OLS<br>(Baseline) | 135 | 83.7 | 126 | 258 |
| EN<br>(Baseline) | 133 | 37.3 | 52.4 | 34.9 |
| LCEN<br>(Baseline) | 33.6 | 12.7 | 18.5 | 33.3 |
| MLP<br>(Baseline) | **20.4** | **7.96** | **11.8** | **29.8** |
| Mechanistic + MLP<br>(This Work) | 31.9 | 8.74 | 14.1 | 34.2 |
| Train Mean<br>(From [30]) | 25.1 | 11.1 | 19.1 | 39.0 |

Mechanistic + MLP residual hybrid model also consistently made predictions with lower errors than the pure mechanistic model, reducing its errors by 1.2-fold on average. Surprisingly, the residual hybrid model was not as good as a pure MLP model in this dataset. We hypothesize this difference occurs because the mechanistic model for this dataset is not as accurate as the one in Ref. [29], because there are additional data available for training, and potentially because the different features and culture settings are more challenging for models that include mechanistic parts (including the pure mechanistic model). These results again confirm the ability of our AutoML method to train strong-performing MLPs, as both models that included MLPs surpassed the other architectures.

### 3.2 Residual hybrid models also improve other important predictions for antibody production

Although predicting the distribution of N-glycans is an important task, there are also other values and metrics of interest in the production of antibodies. The dataset of Ref. [31] comprises antibody titer and galactosylation index measurements for CHO-cell produced antibodies under different feed conditions (FS1–7). Ref. [31] also trained a mechanistic model based on differential equations, named “Mechanistic” in this work. Their model had medium-low errors for half of the titer and most of the galactosylation index predictions (Table 3, top half). However, these errors are train-set errors, as Ref. [31] used all of their data points to train their model, so comparisons with independent test points are not available. We separated points FS2, 3, and 7 to form a test set; the first two because they are the out-of-specification points [31], and FS7 because it was the point for which the mechanistic model of Ref. [31] had the highest prediction errors. As per Section 2.2, we trained data-driven models to expand the models serving as a baseline and an MLP-based residual hybrid model. For all architectures, one set of models for the titer prediction task and another for the galactosylation index task were trained. Due to the low number of training samples (4) and higher number of features (8) than samples, the OLS, EN, and LCEN models had overfit results (despite the use of regularization) when predicting antibody titers, with zero error in all training-set predicitions, but higher errors than the mechanistic model for the test-set predictions.^1^ The MLP model performed considerably better and even surpassed the Mechanistic model on all test-set predictions (Table 3, top half). Finally, the residual hybrid model (again, the mechanistic model followed by an MLP) achieved the best overall performance, surpassing both the “Mechanistic” model of Ref. [31] and the purely data-driven MLPs. The residual hybrid model reduces the average test-set error of the mechanistic model by 1.8 fold on the antibody titer prediction task (Table 3, top half).

**Table 3:** Test-set percent relative errors (PRE) for different model architectures predicting titers (top table) or galactosylation indices (bottom table) for each feed strategy on the dataset of Ref. [31]. Model “Mechanistic” is from Ref. [31]; its PREs are obtained from the published data within. Note that all data points, including FS2, 3, and 7, were used to train the “Mechanistic” model in Ref. [31], so the PREs are train-set values for this model only. Models “OLS”, “EN”, “LCEN”, and “MLP” are data-driven models from this work used as baselines. Model “Mechanistic + MLP” is a residual hybrid model from this work. “Train Mean” is the mean of the training data. The lowest mean test PREs are highlighted in bold.

| Architecture Name | Train set |  |  |  | Test set |  |  |  |
| --- | --- | --- | --- | --- | --- | --- | --- | --- |
|  | FS1 | FS4 | FS5 | FS6 | FS2 | FS3 | FS7 | Mean test PRE |
| Mechanistic<br>(From [31]) | 10.1 | 5.0 | 11.1 | 3.1 | 14.6 | 1.9 | 18.7 | 11.7 |
| OLS<br>(Baseline) | 0 | 0 | 0 | 0 | 12.3 | 11.0 | 25.1 | 16.1 |
| EN<br>(Baseline) | 0 | 0 | 0 | 0 | 11.3 | 7.3 | 24.9 | 14.5 |
| LCEN<br>(Baseline) | 0 | 0 | 0 | 0 | 18.8 | 5.9 | 29.5 | 18.1 |
| MLP<br>(Baseline) | 1.2 | 0.5 | 0.1 | 0.1 | 4.7 | 0.4 | 14.9 | 6.7 |
| Mechanistic + MLP<br>(This Work) | 0.3 | 0.3 | 0.2 | 0.2 | 3.5 | 0.4 | 15.3 | **6.4** |
| Train Mean<br>(From [31]) | 4.1 | 2.5 | 14.5 | 5.6 | 22.6 | 11.4 | 25.7 | 19.9 |
|  | FS1 | FS4 | FS5 | FS6 | FS2 | FS3 | FS7 | Mean test PRE |
| Mechanistic<br>(From [31]) | 10.3 | 14.2 | 5.2 | 17.0 | 10.5 | 10.8 | 29.0 | 16.8 |
| OLS<br>(Baseline) | 0 | 0 | 0 | 0 | 49.1 | 60.2 | 30.3 | 46.5 |
| EN<br>(Baseline) | 23.0 | 5.3 | 3.4 | 7.5 | 52.7 | 53.0 | 25.6 | 43.8 |
| LCEN<br>(Baseline) | 2.1 | 0.7 | 0.1 | 2.0 | 61.4 | 56.5 | 22.7 | 46.9 |
| MLP<br>(Baseline) | 3.9 | 1.1 | 1.9 | 1.5 | 28.8 | 17.3 | 30.0 | 25.4 |
| Mechanistic + MLP<br>(This Work) | 1.5 | 0.4 | 0.7 | 0.8 | 11.4 | 1.1 | 15.7 | **9.4** |
| Train Mean<br>(From [31]) | 29.9 | 7.9 | 0.3 | 12.4 | 47.2 | 29.9 | 33.9 | 37.0 |

On the galactosylation index prediction task, the overfitting issue was less present, affecting only the OLS model. Nevertheless, this task was more challenging, as all data-driven models had a lower performance than the mechanistic model (Table 3, bottom half). As before, the MLP architecture achieved the best results, but its test-set errors were still higher than those of the mechanistic model. Only the residual hybrid model returns more accurate predictions than the mechanistic model, and it does so for all data points (Table 3, bottom half). On average, the residual hybrid model also reduces the test-set error of the mechanistic model by 1.8-fold on the galactosylation index prediction task. These two results further corroborate the potential of residual hybrid models for tasks beyond predicting the distribution of N-glycans, and highlight the ability of our AutoML software to train powerful MLPs even in a data-scarce setting.

The final task investigated involves predicting the concentration of nutrients and metabolites in the medium used to cultivate CHO cells. The dataset of Ref. [32] comprises the levels of 19 chemicals and the amount of biomass in the medium over a culture period of 215 hours. The chemicals include multiple amino acids, sugars, and ammonium. In addition to the large amounts of data gathered by Ref. [32], this dataset is also distinct because each measurement of interest was collected over multiple time points, allowing time series modeling to be done. Ref. [32] trained a flux-based kinetic mechanistic model (named “Mechanistic” in this work) consisting of 103 chemical reaction and transport equations (Table 1 of that work). This mechanistic model had varying performance depending on the prediction task. For example, it performed very well when predicting levels of glucose, lactate, valine, or isoleucine; however, it performed poorly when predicting levels of ammonium, alanine, and biomass. We trained data-driven and residual hybrid models based on the LCEN and RNN architectures to predict levels of lactate, ammonium, biomass, glutamate, aspartate, and asparigine. These dynamic models were trained to output next-hour levels ^1^ based on the levels 1–5 hours prior, with the cutoff depending on the architecture and task. All data-driven and residual hybrid models had more accurate test-set predictions than the mechanistic model (Table 4). In addition, the residual hybrid models trained with a given architecture surpassed the pure data-driven model with the same architecture in all but one case. Overall, the best residual hybrid architectures reduced the test-set prediction error by 736-fold on average (Table 4) and were able to follow the experimental measurements with significant accuracy for both the training and testing periods (Fig. 2). These tests further validate the capabilities of residual hybrid models in yet another context and even despite the fact that the mechanistic model they were based on had limited performance for some metabolites.

**Table 4:** Average test-set percent relative errors (PRE) for different model architectures for predicting metabolite and biomass levels on the dataset of Ref. [32]. Model “Mechanistic” is from Ref. [32]; its PREs are obtained from Fig. 6 of that work. Models “LCEN” and “RNN” are data-driven models from this work used as baselines. Models “Mechanistic + LCEN” and “Mechanistic + RNN” are residual hybrid models from this work. “Train Mean” is the mean of the training data. The lowest PREs are highlighted in bold.

| Architecture Name | Lac | Ammonium | Biomass | Glu | Asn |
| --- | --- | --- | --- | --- | --- |
| Mechanistic<br>(From [32]) | 3.87 | 66.1 | 35.6 | 5.71 | 99.0 |
| LCEN<br>(Baseline) | **0.07** | 0.17 | 1.15 | 3.14 | 13.6 |
| RNN<br>(Baseline) | 1.06 | 1.68 | 2.44 | 0.77 | 1.77 |
| Mechanistic + LCEN<br>(This Work) | 0.36 | **0.02** | **0.23** | 3.01 | 10.4 |
| Mechanistic + RNN<br>(This Work) | 0.34 | 0.20 | 1.98 | **0.25** | **0.53** |
| Train Mean<br>(From [32]) | 9.88 | 44.2 | 60.9 | 20.5 | 591 |

**Figure 2:**
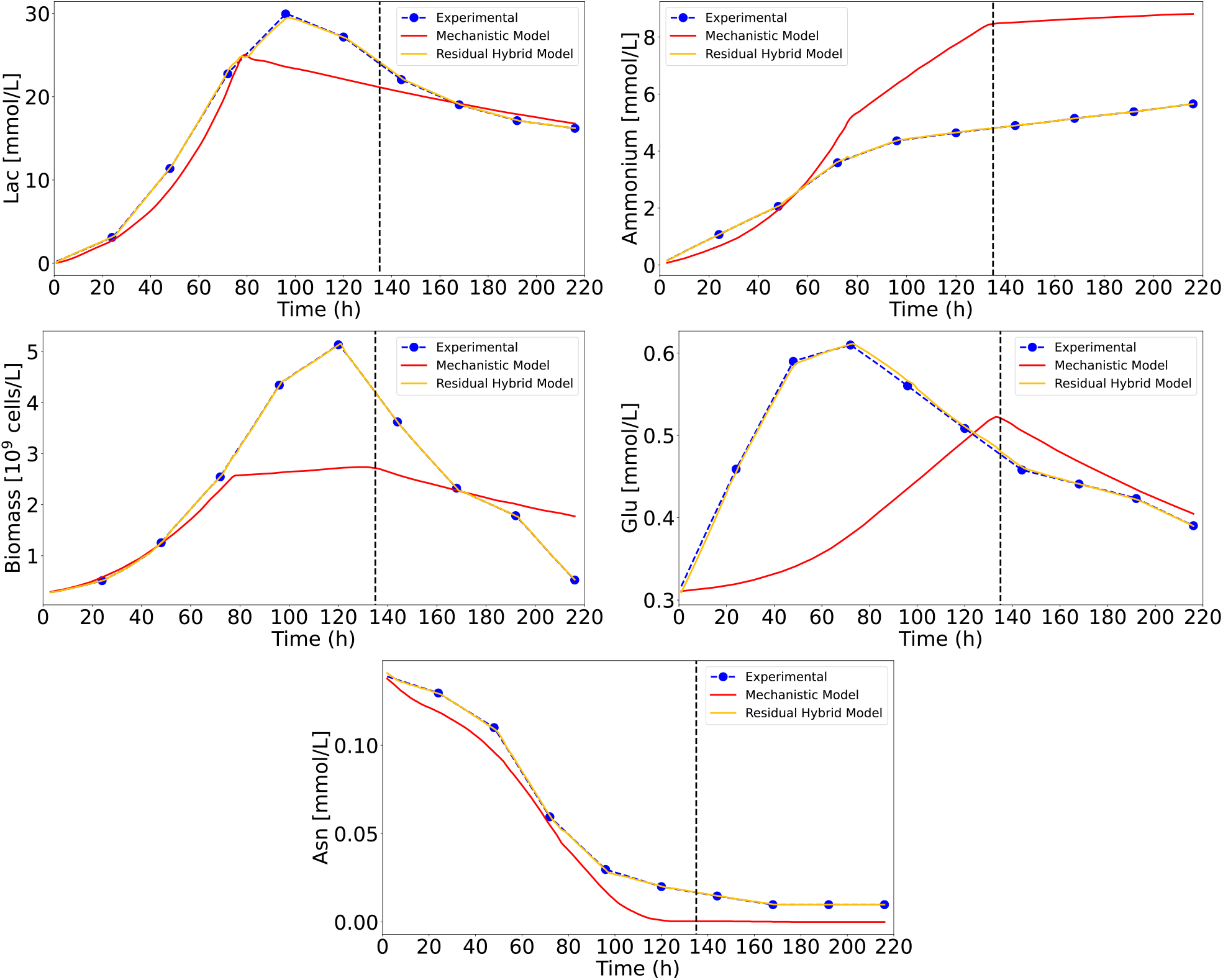
Experimental values and predictions for the mechanistic model (from Ref. [32]) and the best residual hybrid model (“Mechanistic + LCEN” or “Mechanistic + RNN”, this work) for selected metabolites from the Ref. [32] batch culture dataset. The vertical black dotted line separates the training period (t ≤ 135 h) from the test period.

## 4 Discussion

This study constructs residual hybrid models from literature data on the distribution of N-glycans, properties relevant for antibody production, and concentrations of metabolites in CHO cell culture. These datasets not only were built for different tasks (but all relevant for the production of biopharmaceuticals) but also consist of different features, including culture conditions and even a purely autoregressive dataset. The hybrid models combine data-driven models trained in this work and mechanistic models from the literature. As a comparative baseline, purely data-driven models are also trained and tested on the same datasets. The residual hybrid models significantly and consistently had higher prediction accuracy over the mechanistic models, and outperformed the data-driven models in most tasks. Among the four datasets used in this work, residual hybrid models reduce the test-set prediction error of the corresponding mechanistic models by 149-fold on average and always lead to reductions in test-set error for all the predicted variables in the datasets.

The first two datasets (from Refs. [29] and [30]) involve predicting the distribution of (major) N-glycans based on the viable cell density, levels of metabolites (such as Mn and Gal) in the culture, and pH (Section 3.1). Two models were trained by Ref. [29] on their dataset: a mechanistic and an RSM model. The mechanistic model obtained much better results on that dataset. As a baseline, other data-driven models were trained by us. Only an MLP model was able to surpass the mechanistic model, and only for one of the four N-glycans (Table 1). On the other hand, a residual hybrid model consisting of the mechanistic model of Ref. [29] followed by an MLP trained by us was able to reduce the relative errors by 2.45-fold when compared to the mechanistic model, and reductions in prediction errors occurred for all N-glycans (Table 1). Ref. [30] trained a mechanistic model on another dataset for the same task, which we also use to further validate the ability of residual hybrid models to lower prediction errors.

The mechanistic model of Ref. [30] had higher relative errors (which may be explained by their dataset’s being more challenging for prediction than that of Ref. [29]), but it was still relatively accurate. In this task, some data-driven models were able to surpass the mechanistic model: an LCEN model was better than the mechanistic model on 2/4 N-glycans, and an MLP model surpassed the mechanistic model on all N-glycans (Table 2). Finally, a residual hybrid model, which was built in the same manner as that used for the Ref. [29] dataset, was also able to surpass the mechanistic model on all N-glycans, reducing the relative prediction errors by 1.2-fold on average (Table 2). For the dataset of Ref. [30] only, a pure data-driven MLP led to greater reduction in prediction errors than the Mechanistic + MLP residual hybrid model. We hypothesize that this difference in performance may be due to the lower accuracy of the mechanistic model in this task, because there are additional data for training (relative to the dataset of Ref. [29], for example), and because the different features and culture settings in the dataset of Ref. may be more challenging for models that include mechanistic parts (including the pure mechanistic model).

To highlight how residual hybrid models are widely applicable, models were then trained on datasets that contained other relevant metrics for the production of antibodies. Beyond the distribution of N-glycans, this includes the titer, indices that serve as a proxy for N-glycosylation, and the levels of metabolites in the culture. Ref. [31] trained mechanistic models on seven different feed conditions to predict antibody titers and the galactosylation index of the antibodies. Their mechanistic model had medium-low errors for about half of these 14 predictions; however, all of the data points were used to train that mechanistic model, so the errors are biased downwards. Again, data-driven models were trained by us as a baseline; however, most of these had issues with overfitting, so they performed poorly. A notable exception was the MLP model trained to predict titers, which had a test-set error 1.7-fold lower than that of the mechanistic model (Table 3). Residual hybrid models surpassed all of the mechanistic and data-driven models on these tasks, leading to a 1.8-fold reduction in test-set errors when predicting both titers and galactosylation indices. Residual hybrid models led to reduction in prediction errors for every sample in this dataset (Table 3).

Finally, a fourth dataset comprised of the levels of important metabolites over time was used for modeling. Ref. [32] trained a flux-based kinetic mechanistic model on these data. The model achieved mixed success: it was very accurate for some metabolites, but inaccurate for others. As the data of Ref. [32] were time series, LCEN models with *lag* > 0 and RNN models were trained by us both as purely data-driven models and as residual hybrid models. For all five metabolites tested, the data-driven and residual hybrid models surpassed the mechanistic model of Ref. [32] (Table 4). Furthermore, in all but one case, the residual hybrid model containing a given architecture surpassed the data-driven model of the same architecture. On average, the residual hybrid models led to a 736-fold reduction in test-set prediction error (Table 4) and were able to follow the experimental measurements with significant accuracy for both the training and testing periods (Fig. 2).

Overall, the experiments done in this work attest to the high potential of residual hybrid models to substantially reduce the errors of mechanistic models in a variety of tasks, and the high capabilities of our AutoML software to train accurate data-driven and residual hybrid models. The AutoML software used in this work is publicly available, allowing other researchers to reproduce this work and reuse or improve the code for other tasks and datasets. The software is simple to install and use, allowing even non-specialists in data-driven or residual hybrid models to train and use powerful models for any predictive tasks, including those not related to N-glycosylation or antibody production. We provide instructions in Section S1 and on our GitHub (https://github.com/PedroSeber/SmartProcessAnalytics).

## Data availability statement

The data are available in Refs. [29]–[32]. The code used to process the data and the automatic machine learning software used in this article are available in a GitHub repository at https://github.com/PedroSeber/SmartProcessAnalytics.

## Author Contributions

**Pedro Seber**: conceptualization, methodology, software, validation, formal analysis, data curation, writing – original draft, writing – review & editing, visualization.

**Richard D. Braatz**: resources, writing – original draft, writing – review & editing, supervision, project administration, funding acquisition.

## Acknowledgements

This study is supported by the U.S. Food and Drug Administration, Contract No. 75F40121C00090. Any opinions, findings, conclusions, or recommendations expressed in this material are those of the authors and do not necessarily reflect the views of the financial sponsor. The authors would like to thank Dr. Thomas Villiger for providing the raw data of Refs. [29] and [30], which were used to train and test the models in Section 3.1 of this work.

## Supplemental Information

### S1 Using our AutoML tool to train new residual hybrid models

A setup file defining the specific packages and version numbers used in this work is available as `setup.py` on our GitHub (https://github.com/PedroSeber/SmartProcessAnalytics). Instructions on using the AutoML software are also available in the README file and the Examples folder.

### S2 List of hyperparameters used in this work

All possible combinations of the hyperparameters below were cross-validated.

1. For the elastic net (EN) models: *α* = 0 and 20 log-spaced values between −4.3 and 0 (as per `np.logspace(−4.3, 0, 20)`) and L1 ratios equal to [0, 0.1, 0.2, 0.3, 0.4, 0.5, 0.6, 0.7, 0.8, 0.9, 0.95, 0.97, 0.99] were used.
2. For the LCEN models: *α* and L1 ratios as above. *degree* values equal to [1, 2, 3], *cutoff* values between 0 and 4*×* 10^−1^, and *lag* = 0 except for the Ref. [32] dataset, which used *lag* between 1 and 5.
3. For the ANN models: The MLP hidden layer sizes varied for each dataset, with typical sizes varying from 20 to 120 neurons. One to three hidden layers were used. For the Ref. [32] dataset, an LSTM cell with 30–60 neurons before the MLP and a lag between 1 and 4 were also used. Learning rates = [0.05, 0.1, 0.5], batch sizes = 32, the ReLU activation function, weight decays = [0.1, 0.5, 1], 40 epochs, and a cosine scheduler with a minimum learning rate equal to 1/16 of the original learning rate with 10 epochs of warm-up were also used.

## Footnotes

1 Not to be confused with the residual connections (also called skip connections) found in some deep learning models.

2 Except for the procedure for Ref. [32], which uses time series CV due to the nature of its dataset.

3 Keep in mind that the errors for the mechanistic model are all training-set errors, so they are biased downwards.

4 Predicting further in the future without major increases in error is simple; see Table 6 of Ref. [28] for example.

## REFERENCES

[1] B. Imperiali and S. E. O’Connor, “Effect of N-linked glycosylation on glycopeptide and glycoprotein structure,” Current Opinion in Chemical Biology, vol. 3, no. 6, pp. 643–649, 1999.

[2] M. C. Patterson, “Metabolic mimics: The disorders of N-linked glycosylation,” Seminars in Pediatric Neurology, vol. 12, no. 3, pp. 144–151, 2005.

[3] K. T. Schjoldager, Y. Narimatsu, H. J. Joshi, and H. Clausen, “Global view of human protein glycosylation pathways and functions,” Nature Reviews Molecular Cell Biology, vol. 21, no. 12, pp. 729–749, 2020.

[4] S. R. Stowell, T. Ju, and R. D. Cummings, “Protein glycosylation in cancer,” Annual Review of Pathology: Mechanisms of Disease, vol. 10, no. 1, pp. 473–510, 2015.

[5] A. H. Bhat, S. Maity, K. Giri, and K. Ambatipudi, “Protein glycosylation: Sweet or bitter for bacterial pathogens?,” Critical Reviews in Microbiology, vol. 45, no. 1, pp. 82–102, 2019.

[6] J. Jaeken, “Chapter 179 – congenital disorders of glycosylation,” in Pediatric Neurology Part III (O. Dulac, M. Lassonde, and H. B. Sarnat, eds.), vol. 113 of Handbook of Clinical Neurology, pp. 1737–1743, Amsterdam: Elsevier, 2013.

[7] A. Almeida and D. Kolarich, “The promise of protein glycosylation for personalised medicine,” Biochimica et Biophysica Acta (BBA) – General Subjects, vol. 1860, no. 8, pp. 1583–1595, 2016. Glycans in personalised medicine.

[8] W.-L. Ho, W.-M. Hsu, M.-C. Huang, K. Kadomatsu, and A. Nakagawara, “Protein glycosylation in cancers and its potential therapeutic applications in neuroblastoma,” Journal of Hematology & Oncology, vol. 9, no. 1, p. 100, 2016.

[9] M. Ahmed and N.-K. V. Cheung, “Engineering anti-GD2 monoclonal antibodies for cancer immunotherapy,” FEBS Letters, vol. 588, no. 2, pp. 288–297, 2014.

[10] L. Van Landuyt, C. Lonigro, L. Meuris, and N. Callewaert, “Customized protein glycosylation to improve biopharmaceutical function and targeting,” Current Opinion in Biotechnology, vol. 60, pp. 17–28, 2019.

[11] V. Padler-Karavani, H. Yu, H. Cao, H. Chokhawala, F. Karp, N. Varki, X. Chen, and A. Varki, “Diversity in specificity, abundance, and composition of anti-Neu5Gc antibodies in normal humans: Potential implications for disease,” Glycobiology, vol. 18, no. 10, pp. 818–830, 2008.

[12] C. H. Hokke, A. A. Bergwerff, G. W. K. Dedem, J. P. Kamerling, and J. F. G. Vliegenthart, “Structural analysis of the sialylated N- and O-linked carbohydrate chains of recombinant human erythropoietin expressed in Chinese hamster ovary cells. Sialylation patterns and branch location of dimeric N-acetyllactosamine units,” European Journal of Biochemistry, vol. 228, no. 3, pp. 981– 1008, 1995.

[13] K. Bork, W. Reutter, W. Weidemann, and R. Horstkorte, “Enhanced sialylation of EPO by overexpression of UDP-GlcNAc 2-epimerase/ManAc kinase containing a sialuria mutation in CHO cells,” FEBS Letters, vol. 581, no. 22, pp. 4195–4198, 2007.

[14] J. Willard, X. Jia, S. Xu, M. Steinbach, and V. Kumar, “Integrating scientific knowledge with machine learning for engineering and environmental systems,” ACM Comput. Surv., vol. 55, no. 4, 2022.

[15] L. Shen, D. J. Jacob, M. Santillana, X. Wang, and W. Chen, “An adaptive method for speeding up the numerical integration of chemical mechanisms in atmospheric chemistry models: application to GEOS-Chem version 12.0.0,” Geoscientific Model Development, vol. 13, no. 5, pp. 2475–2486, 2020.

[16] A. Derbalah, H. Al-Sallami, C. Hasegawa, A. Gulati, and S. B. Duffull, “A framework for simplification of quantitative systems pharmacology models in clinical pharmacology,” British Journal of Clinical Pharmacology, vol. 88, no. 4, pp. 1430–1440, 2022.

[17] G. Cybenko, “Approximation by superpositions of a sigmoidal function,” Mathematics of Control, Signals, and Systems, vol. 2, pp. 303–314, 1989.

[18] K. Hornik, “Approximation capabilities of multilayer feedforward networks,” Neural Networks, vol. 4, no. 2, pp. 251–257, 1991.

[19] C. Rudin, “Stop explaining black box machine learning models for high stakes decisions and use interpretable models instead,” Nature Machine Intelligence, vol. 1, pp. 206–215, 2019.

[20] J. Štor, D. E. Ruckerbauer, D. Széliová, J. Zanghellini, and N. Borth, “Towards rational glycoengineering in CHO: from data to predictive models,” Current Opinion in Biotechnology, vol. 71, pp. 9–17, 2021.

[21] C. Kontoravdi and I. Jimenez del Val, “Computational tools for predicting and controlling the glycosylation of biopharmaceuticals,” Current Opinion in Chemical Engineering, vol. 22, pp. 89–97, 2018.

[22] C. Liang, A. W. Chiang, A. H. Hansen, J. Arnsdorf, S. Schoffelen, J. T. Sorrentino, B. P. Kellman, B. Bao, B. G. Voldborg, and N. E. Lewis, “A Markov model of glycosylation elucidates isozyme specificity and glycosyltransferase interactions for glycoengineering,” Current Research in Biotechnology, vol. 2, pp. 22–36, 2020.

[23] P. Seber and R. D. Braatz, “Linear and neural network models for predicting N-glycosylation in Chinese Hamster Ovary cells based on B4GALT levels,” bioRxiv, 2023.

[24] M. Aykol, C. B. Gopal, A. Anapolsky, P. K. Herring, B. van Vlijmen, M. D. Berliner, M. Z. Bazant, R. D. Braatz, W. C. Chueh, and B. D. Storey, “Perspective—combining physics and machine learning to predict battery lifetime,” Journal of The Electrochemical Society, vol. 168, no. 3, p. 030525, 2021.

[25] L. Liao and F. Köttig, “Review of hybrid prognostics approaches for remaining useful life prediction of engineered systems, and an application to battery life prediction,” IEEE Transactions on Reliability, vol. 63, no. 1, pp. 191–207, 2014.

[26] H.-T. Su, N. Bhat, P. Minderman, and T. McAvoy, “Integrating neural networks with first principles models for dynamic modeling,” IFAC Proceedings Volumes, vol. 25, no. 5, pp. 327–332, 1992.

[27] M. L. Thompson and M. A. Kramer, “Modeling chemical processes using prior knowledge and neural networks,” AIChE Journal, vol. 40, no. 8, pp. 1328–1340, 1994.

[28] P. Seber and R. D. Braatz, “LCEN: A novel feature selection algorithm for nonlinear, interpretable machine learning models,” 2024. 2402.17120.

[29] D. J. Karst, E. Scibona, E. Serra, J.-M. Bielser, J. Souquet, M. Stettler, H. Broly, M. Soos, M. Morbidelli, and T. K. Villiger, “Modulation and modeling of monoclonal antibody N-linked glycosylation in mammalian cell perfusion reactors,” Biotechnology and Bioengineering, vol. 114, no. 9, pp. 1978–1990, 2017.

[30] T. K. Villiger, E. Scibona, M. Stettler, H. Broly, M. Morbidelli, and M. Soos, “Controlling the time evolution of mAb N-linked glycosylation - Part II: Model-based predictions,” Biotechnology Progress, vol. 32, no. 5, pp. 1135–1148, 2016.

[31] P. Kotidis, P. Demis, C. H. Goey, E. Correa, C. McIntosh, S. Trepekli, N. Shah, O. V. Klymenko, and C. Kontoravdi, “Constrained global sensitivity analysis for bioprocess design space identification,” Computers & Chemical Engineering, vol. 125, pp. 558–568, 2019.

[32] M. Kastelic, D. Kopač, U. Novak, and B. Likozar, “Dynamic metabolic network modeling of mammalian Chinese hamster ovary (CHO) cell cultures with continuous phase kinetics transitions,” Biochemical Engineering Journal, vol. 142, pp. 124–134, 2019.

[33] F. Zamorano, A. Vande Wouwer, R. Jungers, and G. Bastin, “Dynamic metabolic models of cho cell cultures through minimal sets of elementary flux modes,” Journal of Biotechnology, vol. 164, no. 3, pp. 409–422, 2013.

[34] F. Pedregosa, G. Varoquaux, A. Gramfort, V. Michel, B. Thirion, O. Grisel, M. Blondel, P. Prettenhofer, R. Weiss, V. Dubourg, J. Vanderplas, A. Passos, D. Cournapeau, M. Brucher, M. Perrot, and E. Duchesnay, “Scikit-learn: Machine learning in Python,” Journal of Machine Learning Research, vol. 12, pp. 2825–2830, 2011.

[35] A. Paszke, S. Gross, F. Massa, A. Lerer, J. Bradbury, G. Chanan, T. Killeen, Z. Lin, N. Gimelshein, L. Antiga, A. Desmaison, A. Kopf, E. Yang, Z. DeVito, M. Raison, A. Tejani, S. Chilamkurthy, B. Steiner, L. Fang, J. Bai, and S. Chintala, “PyTorch: An imperative style, high-performance deep learning library,” in Advances in Neural Information Processing Systems 32, pp. 8024–8035, Curran Associates, Inc., 2019.

